# X-ray Diffraction Reveals Periodicity in Murine Neocortex

**DOI:** 10.1101/2024.10.13.618071

**Authors:** Sasha Murokh, Ezekiel Willerson, Alexander Lazarev, Pavel Lazarev, Joshua C. Brumberg

## Abstract

**Background:** Sensory experience impacts brain development. In the mouse somatosensory cortex, sensory deprivation via whisker trimming induces reductions in the perineuronal net (PNN), the size of neuronal cell bodies, the size and orientation of dendritic arbors, the density of dendritic spines, and the level of myelination, among other effects.

**New Methods:** Here, we measured the X-ray diffraction patterns of mouse brain tissue to establish a novel method for examining nanoscale brain structures. Two groups of mice were examined: a control group and one that underwent 30 days of whisker-trimming from birth - an established method of sensory deprivation that affects the mouse barrel cortex (whisker sensory processing region of the primary somatosensory cortex). Mice were perfused, and primary somatosensory cortices (barrel cortex) were isolated for immunocytochemistry and X-ray diffraction imaging.

**Results:** X-ray images were characterized using a specially developed machine-learning approach, and the clusters that correspond to the two groups are well separated in the space of the principal components. The obtained values for sensitivity/specificity are 1/0.93, and the receiver operator curve classifier is 0.99.

**Conclusions:** We hypothesize that such separation is related to the development of different nanoscale structural components in the brains of control and sensory deprived mice. The effects of these nanoscale structural formations can be seen in PNN and other micro- and macro-scale structures and assemblies.

## 1. Introduction

X-ray absorption has been widely used in medicine since its discovery. However, another effect, X-ray diffraction (or, more generally, scattering), has not been implemented widely. Initially, it was utilized for X-ray crystallography, determining the atom arrangements in crystalline solids (Bragg, 1914). Later, bio-crystallography based on crystallized biological molecules facilitated the revealing of the molecular content of complex structures, including myoglobin, hemoglobin, and DNA; see (Shi, 2014; Higgins & Lea, 2017) for historical overviews. However, the X-ray scattering approach can be applied directly to biological tissues, and if there are periodic structures, they will manifest themselves in characteristic scattering patterns. In this, any periodicity, *d*, of the structure, such as regular intermolecular distances in molecular packing or the pitch in multiple-helix fibrils, will appear as momentum transfer, *q* = 2πn/*d*. In particular, the 64 nm pitch of the triple helix of collagen produces peaks in the X-ray scattering patterns at 0.1 nm^-1^ and its multiples (Worthington & Tomlin, 1955).

Small-angle (SAXS, 0.1 - 5 nm^-1^) and wide-angle (WAXS, 3 - 45 nm^-1^) X-ray scattering techniques have been widely used to characterize neuronal tissues in humans (De Felici, et al., 2008) and animals (Yagi, 2011; Choi, et al., 2017; Dahal, et al., 2020). They can be combined with scanning and tomography methods; see (Omori, et al., 2023) for the description of recent advances in this field. The strongest signal is associated with myelin. In the SAXS region, there are pronounced peaks at about 1 and 1.5 nm^-1^. In the WAXS region, the peak originating from the lipid component of myelin was observed at about 14.5 nm^-1^, while the aqueous component produced a peak at about 20 nm^-1^ (De Felici, et al., 2008). The former peak is much less pronounced in the gray matter than in the white matter of the brain tissues. There is also a broad peak at about 6 nm^-1^, which was attributed to myelin protein (Inouye & Kirschner, 1984). Variations of the X-ray diffraction signal were studied in relation to Alzheimer’s Disease (Dahal, et al., 2020), where the effect of the amyloid burden on SAXS was examined, and Parkinson’s disease (Carboni, et al., 2017), where the XRD tissue scans exhibited increased amounts of crystallized cholesterol. The modifications in the myelin XRD signal caused by the brain tumor were also observed (Siu, et al., 2005; Jensen, et al., 2011).

In the present paper, we use the WAXS measurements to examine the nanoscale structures associated with sensorial deprivation in mice achieved by trimming whiskers. It has previously been shown that sensory deprivation induced by whisker trimming alters the animal’s physiology (Simons & Land, 1987) and behavior (Carvell & Simons, 1996, reviewed in Chen & Brumberg, 2021). At the anatomical level, the overall number of cells within the barrel cortex is not affected by trimming (McRae et al. 2007, Barrera et al. 2013), but the density of the perineuronal net (PNN, McRae et al., 2007), dendritic architecture (Chen et al. 2012), the density of dendritic spines (Chen et al. 2015), axonal myelination (Barrera et al. 2013), and microglia (Kalambogias et al. 2021) are all drastically impacted. Given the ease and reliability of staining we will utilize PNN staining through the use of wisteria floribunda (WFA) histochemical staining to confirm that sensory deprivation is having its intended effect. What is occurring at the nanoscale is unknown and is the focus of the present paper.

We compare the X-ray diffraction patterns of brain tissue of mice with trimmed whiskers to brain tissue of control mice and observe that after standard azimuthal integration, the scattering profiles are indistinguishable. However, such integration eliminates all effects related to the anisotropy of scattering. To overcome this deficiency, we develop a novel approach based on two-dimensional Fourier transformation and apply a machine-learning procedure complemented by the Principal Component Analysis. The obtained metrics demonstrate an excellent cluster separation, indicating the formation of different nanoscale structures for the two groups of mice. We also perform histological studies of PNNs as a benchmark to ensure the consistency of the whiskers-trimming procedure and confirm the biological impact of trimming.

## 2. Methods

### 2.1. Animals and Trimming procedure

Pregnant CD-1 mouse dams were ordered from Charles River to give birth in-facility. Mice were housed as dams plus pups for the duration of the experiment, with each litter serving as either the control or experimental group separately. Animals were housed in standard mouse cages and received *ad libitum* access to food and water. Experiments were in accordance with NIH standards for the use of animals for biomedical research and approved by the Queens College, City University of New York, Institutional Animal Care and Use Committee (protocol #208).

Sensory deprivation via whisker-trimming took place every other day from postnatal day (P) 0 (day of birth) to P29. For the experimental “trimmed whiskers” (TW) group, pups were briefly removed from their litter, and whiskers were trimmed bilateral using Vanna scissors (F.S.T., Catalog #’s 15002-08 or 15018-10). Control group (“untrimmed”) pups were similarly removed from their litter, handled identically, and then returned to their home cage without trimming. From P0-P12, whisker-trimming and handling were performed on awake pups. From P13-P29, trimming and handling were done under light anesthesia to ensure proper trimming and reduce the risk of injury to pups. Isoflurane (3% in room air) was administered for 30 seconds to 1 minute prior to trimming, resulting in the pups being lightly unconscious. Untrimmed control pups were also subjected to isoflurane. Pups were returned to their litter as they regained consciousness.

Mice were euthanized with an intraperitoneal injection (Euthasol, 100mg/kg) and perfused on P30. Deep anesthesia was confirmed when a toe pinch no longer elicited a paw withdrawal. Mice were perfused with 0.9% NaCl followed by 4% paraformaldehyde dissolved in 0.1M phosphate-buffered saline (PBS). Following removal, brain tissue was preserved in 4% paraformaldehyde in 0.1M PBS at 4°C.

### 2.2. Staining for perineuronal nets

For tissue staining, brains were submerged in 4%cparaformaldehyde in 0.1M PBS for 24 hours before being placed in a 30% sucrose solution for three days (or until sunk). The barrel cortex was located using an adult mouse brain atlas (Paxinos and Franklin 2019) and sectioned at 40 μm on a freezing stage microtome (Epredia, HM450). For perineuronal net visualization, tissue sections were washed in 0.01M PBS (3 × 5 minutes) before submerging in a solution of 0.06% hydrogen peroxide combined with 1% methanol by volume in 0.01M PBS to quench endogenous peroxidase activity (30 minutes). Following three washes of 5 minutes in 0.01M PBS, free-floating tissue was further incubated in a solution of biotinylated wisteria floribunda agglutinin (Sigma-Aldrich, L1516-2MG; 1:500 ratio) with 2% normal goat serum dissolved in 0.01M PBS for 72 hours at 4°C. After this initial incubation, tissue was rinsed again three times (5 minutes each) in 0.01M PBS before immersing in an avidin-biotin complex (ABC; Vector labs, PK-4000) in the same concentration PBS for 30 minutes. Following another three washes of 5 minutes each in 0.01M PBS, the perineuronal nets were visualized by reacting the bound ABC with a 3,3′-Diaminobenzidine tetrahydrochloride (Sigma-Aldrich, D5905; 2mg/mL PBS), nickel chloride (0.04%), and hydrogen peroxide (0.15%) solution dissolved in 0.01M PBS. Brain slices were mounted and then allowed to dry on gelatinized glass slides for 24 hours. Gelatinized slides were prepared ahead of time by submerging slides in porcine gelatine (5g/1000mL water) with chromium potassium sulfate (0.5g). Tissue was then dehydrated and defatted by submerging slides in increasing concentrations of ethanol (50%, 70%, 95%, 100%) before finally washing in Safeclear for 5 minutes (Fisher-Scientific, 314-629). Slides were coverslipped with Permount mounting medium (Fisher Chemical, SP15) and allowed to air dry prior to observation.

### 2.3. Sectioning for X-ray diffraction

The tissues used for x-ray diffraction were from different animals and not sunk in sucrose but removed from the brain in the same way as the tissue utilized for immunohistochemistry. Seven animals per group were utilized for x-ray diffraction (Trimmed Whiskers: 3 female, 4 male; Control: 5 female, 2 male). Left and right hemisphere barrel cortices were sectioned (described below) in their entirety for each subject, resulting in 14 samples per group. Using coordinates from the Paxinos and Franklin’s mouse brain atlas (2019), the barrel cortex was determined to have a total length of 2.4 mm along the rostrocaudal axis, spanning the area 3.88 mm to 6.28 mm from the rostral end of the brain (olfactory bulb). This corresponds to approximately 0.37 mm anterior and 2.03 mm posterior to bregma (Paxinos and Franklin’s). Initial cuts were made as closely to these measurements as possible, using 0.25 mm and 0.5 mm brain matrices (Zivic Instruments; Braintree Scientific) and an mm ruler. Mediolaterally, the barrel cortex was determined to span the region 2.25 to 4 mm laterally from the longitudinal fissure. The cortex was sectioned away from the underlying white matter just superior to the corpus callosum, with the cortical plate having a thickness of about 1 mm. The total isolated section size measured approximately 2.4 mm on the rostrocaudal axis, 1.75 mm mediolaterally, and 1 mm thick (isolated cortex). Post-sectioning, tissue was stored in 0.01M PBS at 4°C, and experimenters performing X-ray diffraction were blinded to the condition. The resultant sample preserved the majority of the barrel cortex intact for the subsequent x-ray diffraction studies.

### 2.4. Quantification of Immunohistochemistry

Stained PNNs were quantified using luminance measurements embedded within the Neurolucida software (MBF Bioscience), which averages the brightness of each pixel in a defined region to create a “brightness” score. First, the barrel cortex was traced using the Paxinos and Franklin’s atlas (2019) as a guide. Second, because very few PNNs exist in white matter, white matter local to each barrel cortex was also traced to be used as a comparison point for the barrel cortex. Third, white matter was imaged with lighting adjusted in intensity to normalize images to a 200 (+/-4) brightness score. Fourth, the barrel cortex was imaged at the same light level. Lastly, brightness scores from the barrel cortex were normalized by dividing the luminance values from the underlying white matter to give a final brightness ratio. Thus, a final brightness ratio of “1.0” would represent a barrel cortex with no staining and a higher brightness value, representing fewer PNNs. Four mice were utilized per group, 2 sections per hemisphere, for a total of 16 sections per group.

### 2.5. Diffraction measurements

We used a custom-built transmission-mode diffractometer (Lazarev et al.,2023) comprising a Xenocs source, Genix 3D Cu with Fox 12-53 Cu Mirror (λ = 0.1540562 nm), an Advacam MiniPIX TPX3 detector (256×256 pixel sensor array, with the pixel size of 55 μm^2^), and a sample stage. The image of the device is shown in Fig. 1(a), with its schematics provided in Fig. 1(b). The X-ray beam with a flux of about 10^8^ photons per mm^2^ sec is collimated with an approximate spot size of 0.75 mm in diameter. The samples were placed in a plastic container to prevent drying. The diffractometer was calibrated using a sample of powdered silver behenate. The samples were scanned for 2 minutes at a 13 mm sample-to-detector distance. With such a small distance, the WAXS signal is primarily detected. Measurement data was saved as an array of 256×256 integers. A typical diffraction pattern is shown in Fig. 1(c).

**Figure 1.**
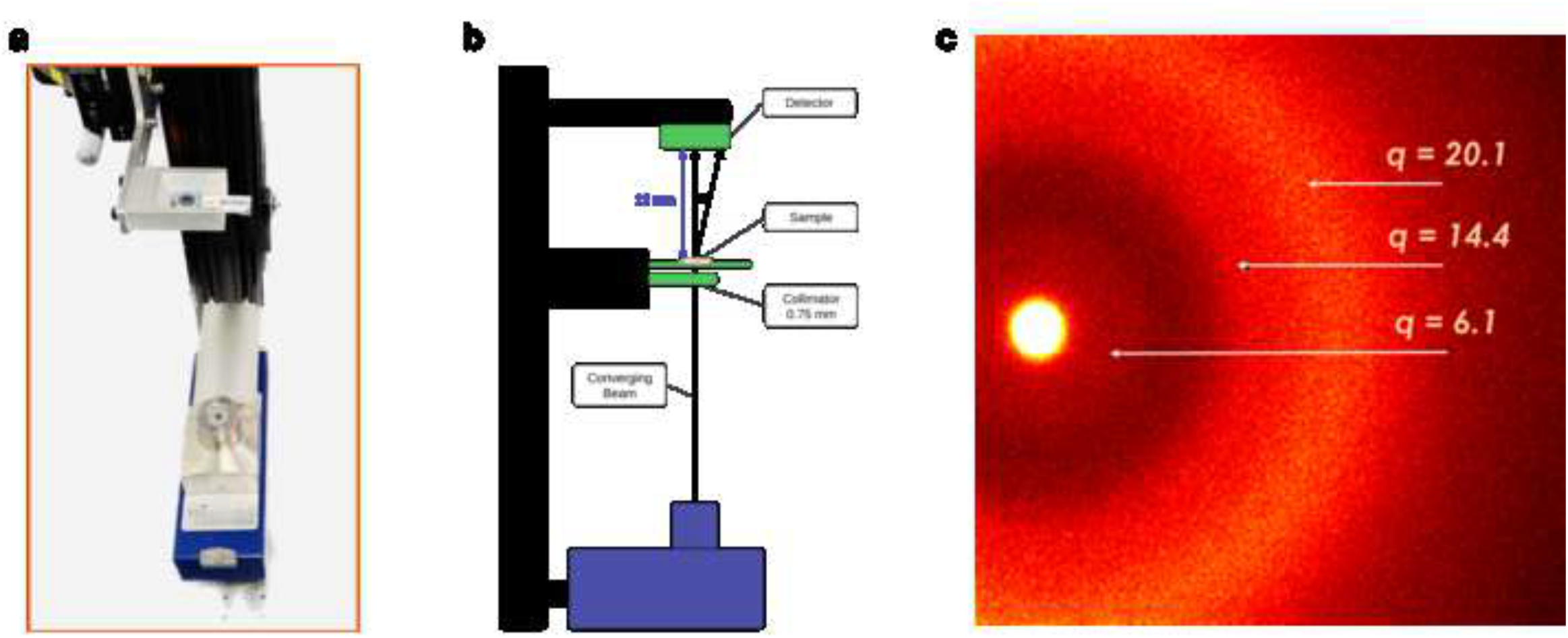
Experimental Setup. (a) Image of the diffractometer. (b) Schematics of the diffractometer. (c) The X-ray diffraction pattern produced by the sample containing a mouse brain slice (2.4 × 1.75 × 1 mm). The bright circle at the center is the unscattered image of the primary beam. The bright concentric rings are associated with constructive interference of light scattered by periodically structured material. The prominent features and corresponding values of the momentum transfer are indicated by arrows.

### 2.6. Data analysis

Each diffraction image is digitally represented as a 256×256 matrix. As the first step, the faulty pixels are removed. Next, azimuthal integration can be performed, and the scattered intensity becomes the function of the distance to the center. This distance can be converted to the momentum transfer, *q*. The concentric rings of Fig. 1(c) appear as specific features in this dependence with momentum transfer values of 6.1, 14.4, and 20.1 inverse nanometers. In simple, straightforward cases, the position and amplitude of these features provide enough information for classifications. However, more advanced data representation and machine-learning methods are needed for complex situations when azimuthal anisotropy plays a significant role.

For this work, we developed an approach based on the two-dimensional Fourier transformation (2DFT) of the image matrix. We use the version of this procedure implemented in the NumPy library. When 2DFT is performed, the 256×256 matrices of the Fourier coefficients (65,536 total) are obtained for each image. To eliminate the contribution of the primary beam, the auxiliary matrices are constructed with zeroes everywhere except the locations of the bright circles. 2DFT of these matrices contain the Fourier coefficients of the primary beam images. The difference between 2DFTs of original and auxiliary matrices provides the Fourier coefficients associated only with the signal scattered from the tissue. Obtained Fourier coefficients are complex numbers, but we use only their magnitudes for further analysis. The elements of the acquired matrices are normalized by the element at (0,0), i.e., brightness, to exclude the irregularities produced by fluctuations in the exposure times and material densities. Next, standard scaling from the scikit-learn library (Pedregosa, et al., 2011) is used to prevent arbitrary pixels and Fourier coefficients from having too much weight.

In the following step, a procedure similar to face recognition (eigenface) (Sirovich & Kirby, 1987; Turk & Pentland, 1991) is accomplished. Each of 28 (256×256) arrays of Fourier coefficients is flattened, i.e., it is represented as a row with 65,536 elements. For the resulting 65,536×28 matrix, a principal component (PC) analysis is performed. Accordingly, each sample is described as a point in the PC space. We applied the Logistic Regression classifier from the scikit-learn library (Pedregosa, et al., 2011) to determine the sensitivity of the group separation (proportion of the trimmed whiskers samples correctly identified), its specificity (proportion of the control samples correctly identified), and the receiver operating characteristics (ROC) curve. The latter illustrates the performance of a binary classifier at varying threshold values. The area under the ROC curve is also used as a metric.

## 3. Results

### 3.1. PNN staining results

To confirm that trimming was having its intended biological impact we analyzed the PNN density in layer IV. In previous work, we found that layer 4 (L4) of the barrel cortex houses the greatest quantity of PNNs (Chu et al., 2018). We also found that whisker-trimming affects PNNs most significantly within L4 of the barrel cortex (Chu 2015; McRae et al., 2007; Uneno et al. 2017). Therefore, we focused our analysis specifically on this layer. Current results confirm previous findings. Whisker-trimming resulted in a significant reduction of L4 barrel cortex PNNs in TW mice when compared with Control mice (see Fig. 2(a,b) for representative samples). The average brightness ratio for Control vs. TW mice was 0.56 and 0.62 respectively, (one-tailed t-test, p=0.025, see Fig. 2(c)). Trimming resulted in an approximate 10% increase in brightness, providing a positive control for the impact of whisker trimming within the somatosensory cortex. Our results are in alignment with several previous studies (Ueno et al. 2017, Chu 2015) but in contrast with McRae et al (2007) which did not see decreases using a different WFA stain, but they did see a decrease in aggrecan (a component of the PNN) utilizing a monoclonal antibody. Our use of WFA staining was to provide a readout that the sensory deprivation paradigm was evoking the exepected biological effect and not to necessarily explain the x-ray difraction results.

**Figure 2.**
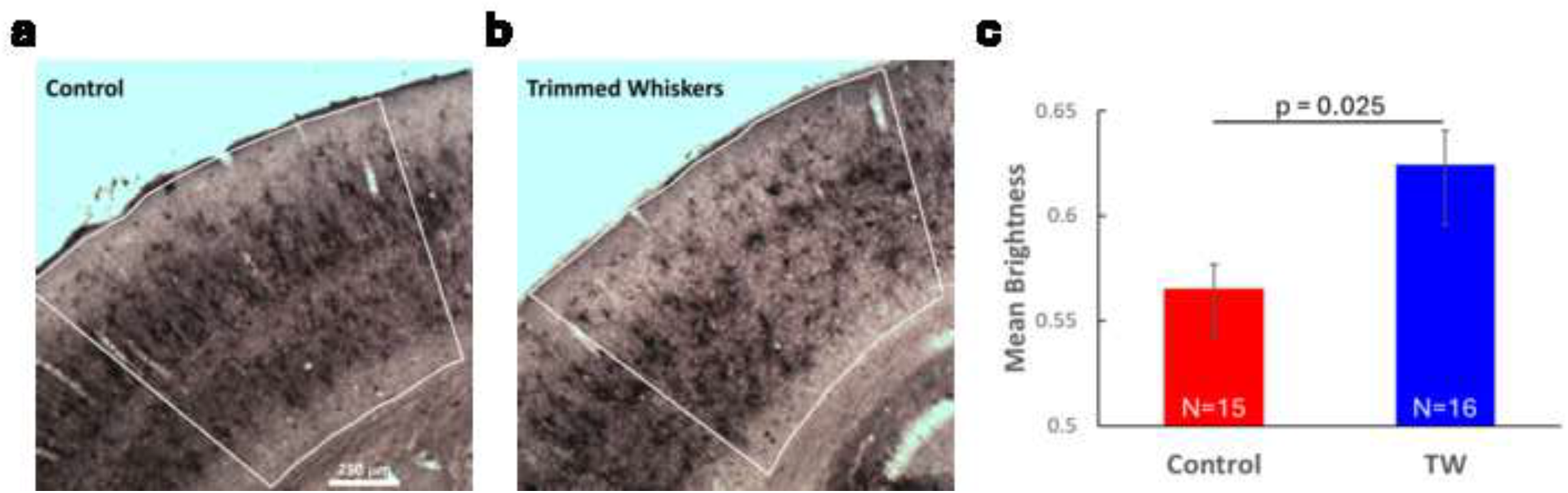
PNN staining. Representative PNN staining of barrel cortex for (a) control and (b) trimmed (TW) mice. Outlines indicate barrel cortex for each sample. (c) Quantification of brightness within layer IV. Mice whose whiskers have been trimmed show higher brightness scores compared with control mice. Bars represent population means and error bars are indicative of one standard error of the mean, N’s are reflective of number of samples.

### 3.2. Analysis of diffraction patterns

To determine if changes in the brain structure at the microscale were correlated with the nanoscale structural changes, we measured the X-ray diffraction patterns for each of the 28 samples. The typical result is shown in figure 1(c), exhibiting several concentric rings. Visually, the patterns of the Control mice and those with trimmed whiskers cannot be distinguished. After azimuthal integration and unit conversion, the intensity can be plotted as a function of the momentum transfer magnitude, figure 3(a). Three features (peaks and shoulders) are identified, approximately at 6.1, 14.4, and 20.1 inverse nanometers. The first shoulder was previously attributed to myelin protein (Inouye & Kirschner, 1984). The second and third features are related to myelin’s lipid and aqueous components (De Felici, et al., 2008). There is also an amorphous scattering signal at small *q*, which obeys Porod’s Law (Porod, 1951), i.e., it is proportional to inverse *q* in the fourth power. The general shape of the curve is similar to what has been published for the gray matter of the brain (De Felici, et al., 2008).

**Figure 3.**
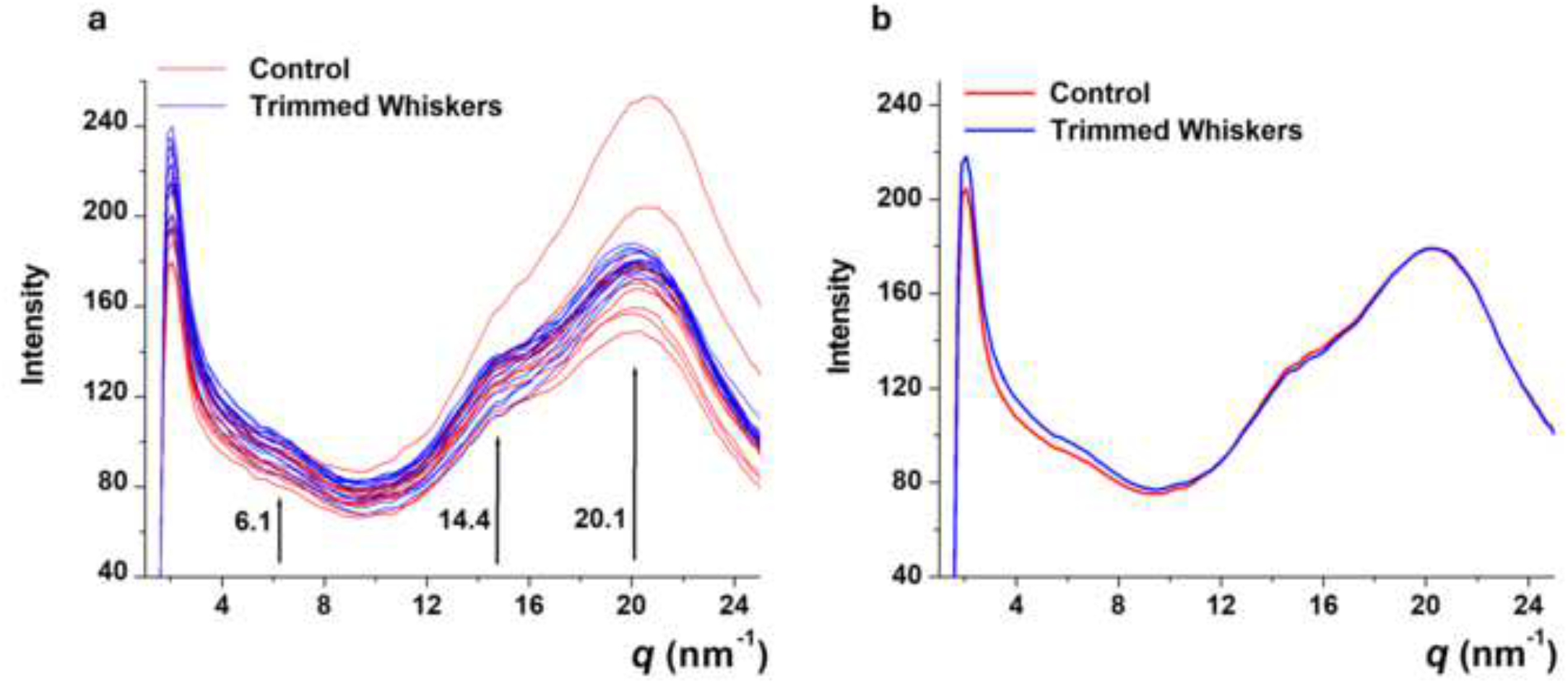
Momentum transfer. (a) Dependencies of the scattered intensity on the momentum transfer magnitude for all measured samples. The prominent features and corresponding values of the momentum transfer are indicated by arrows. (b) Dependencies of the scattered intensity on the momentum transfer magnitude for the mean curves, with the Control and TW shown in red and blue, respectively.

One can see that the Control and TW dependencies are not different, with almost the same positions of the features for all the curves and the magnitudes unrelated to the specifications. As can be further seen in figure 3(b), for the mean curves, the only difference can be seen at small *q* where the amorphous scattering dominates. The stronger signal from the sensory deprived mice supports our conclusion below that these mice do not develop the same brain structure as untrimmed Control mice. We developed and performed the advanced machine-learning-based procedure described in Section 2.6. The explained variances for the determined principal components are shown in Table 1.

**Table 1.**
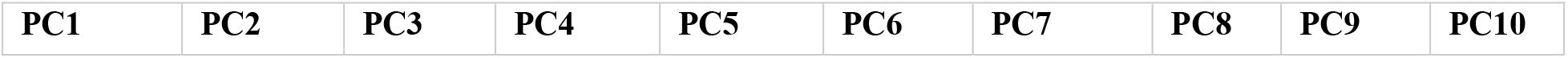
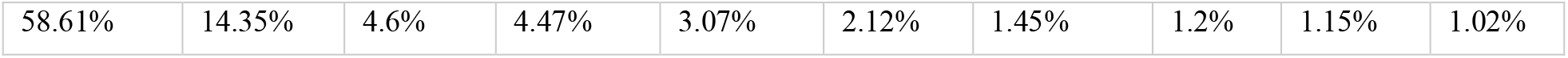
Explained variance (first raw) and cumulative explained variance (second raw) for the first ten dimensions of the principal component analyses (PCA).

| PC1 | PC2 | PC3 | PC4 | PC5 | PC6 | PC7 | PC8 | PC9 | PC10 |
| --- | --- | --- | --- | --- | --- | --- | --- | --- | --- |
| 58.61% | 14.35% | 4.6% | 4.47% | 3.07% | 2.12% | 1.45% | 1.2% | 1.15% | 1.02% |

The results in the 3D PC space are presented in figure 4. While some separation can be seen, it is not enough to obtain good classification metrics. However, the classification becomes excellent when we perform the Logistic Regression procedure in the 10-dimensional PC space. Either sensitivity or specificity has a perfect value of 1, while the other metric is 0.93. The obtained ROC curve is shown in figure 5. The area under the curve is 0.99, which is next to perfect. We interpret this as a formation of different types of structures at the nanoscale in Control and TW mice.

**Figure 4.**
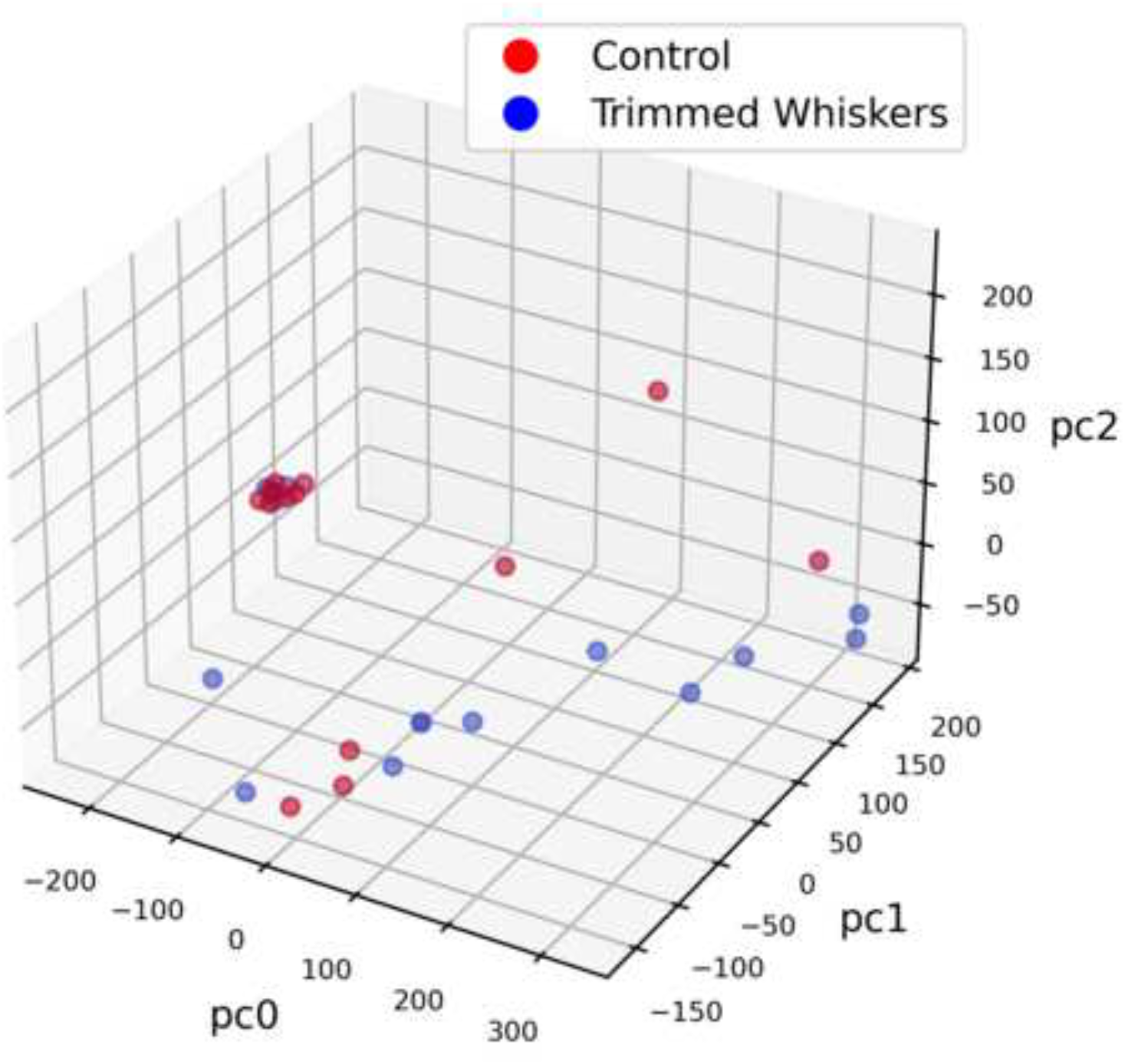
Principal components separate trimmed versus control animals. (a) The PCA-transformed data within the 3D PC space. The red and blue points are associated with the Control mice and those with trimmed whiskers, respectively.

**Figure 5.**
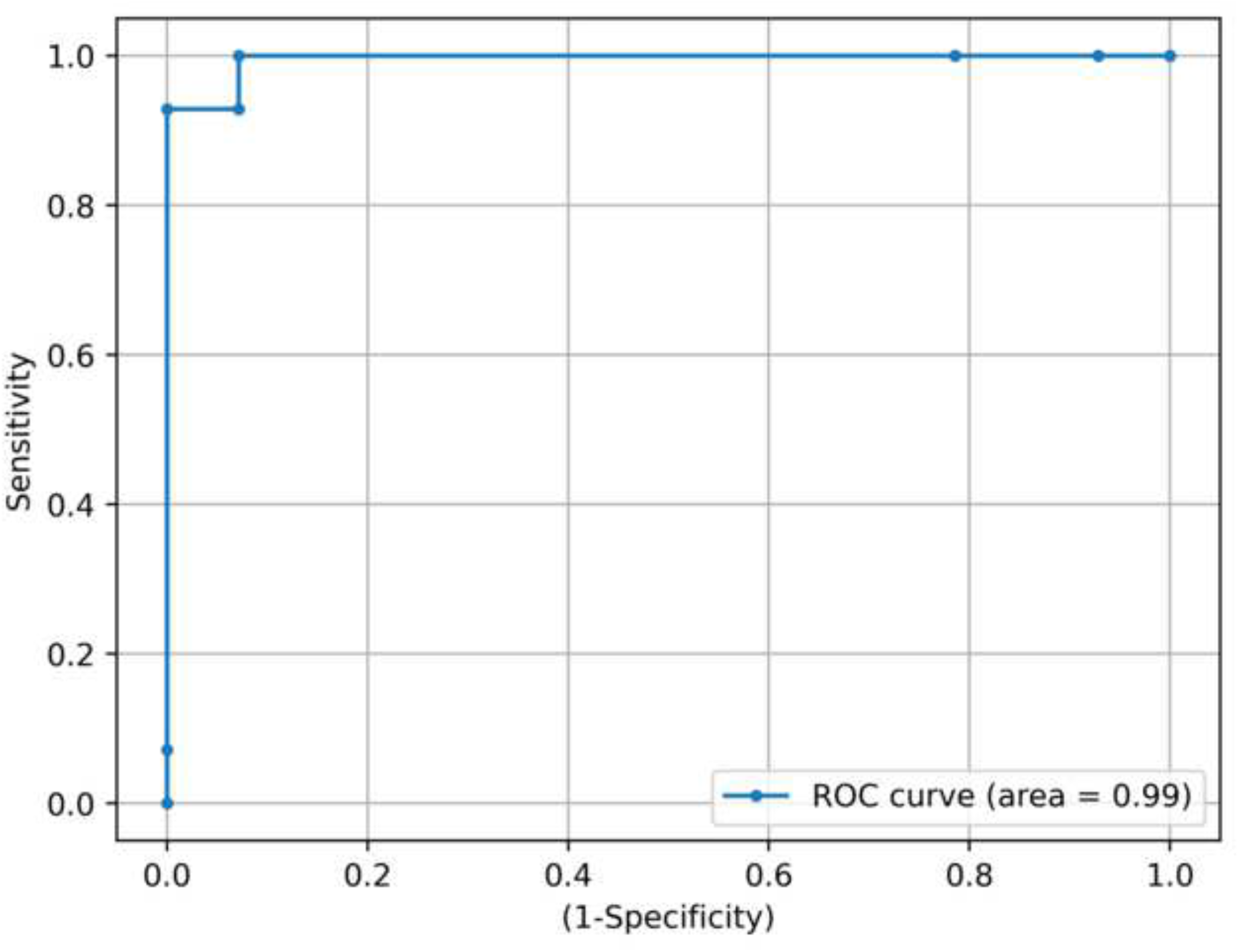
Receiver Operating Characteristic (ROC) curve. The ROC curve shows the sensitivity and specificity of the separation method.

## 4. Discussion

Our current results confirm previous studies that have demonstrated that sensory deprivation via whisker-trimming reduces perineuronal nets in the mouse barrel cortex, as demonstrated by immunohistochemistry (Chu et al. 2018). Here, in a novel approach, we attempt to reveal the origin of these changes on a nanoscale level. We employ the X-ray diffraction (XRD) technique and show that this approach can be used on brain tissue in a novel way to detect structural changes. Isolated barrel cortices from mice that had either undergone 30 days of whisker-trimming or simple handling (controls) were subjected to x-ray diffraction analysis. The characteristic features at momentum transfer values of 6.1, 14.4, and 20.1 inverse nanometers were determined for all samples, with the control and trimmed-whiskers ones being indistinguishable. These results are similar to previous studies where the obtained peaks are related to the myelin protein (Inouye & Kirschner, 1984) and the lipid and aqueous components of the myelin (De Felici, et al., 2008), respectively. However, after a specifically developed novel advanced analysis based on the 2D Fourier transformation of XRD images and the face recognition procedure in the reciprocal space, we reveal the formation of distinct clusters for the control and trimmed-whiskers groups in the principal component space. The Logical Regression classifier for 10 PCs provides a sensitivity or specificity of 1 (with the other metric being 0.93), and the area under the ROC curve is 0.99, which is next to perfect. This demonstrates that the brains of these two groups have different structures at the nanoscale level. We argue that sensorial deprivation leads to the formation of nanoscale structures, different from conventional brain development. However, the exact changes observed in the XRD experiments are still unclear at the present stage. In the context of our analysis, all the principal components are the linear combinations of 65,536 Fourier coefficients. The comparison of the peak positions and magnitudes allows for a clear physical interpretation, but the specific measurements made in our experiment do not allow for the separation of two groups of mice. At the same time, the analysis in terms of two-dimensional Fourier coefficients provides almost perfect separation of the mice groups, but the physical origin remains unclear. The determination of the interconnections between the Fourier coefficients and actual parameters of the nanoscale structures is a long-term project, with our current work being the first step in this direction.

Currently, our methodology describes a technique for examining ex-vivo postmortal tissue to determine nanoscale structural changes related to alternations of micro and macro-scale brain structures. It is feasible to think that this could eventually lead to ex-vivo use for living organisms or even in-vivo, where X-ray diffraction becomes an effective diagnostic tool. In particular, it can be implemented to quickly examine tissues during brain surgeries to determine the extent of the operation. While in-vivo studies for humans are extremely difficult because of the skull size, it might be possible for various animal models. Additional studies are needed to connect our current observations and the actual nanoscale content of brain tissue. A deeper investigation is necessary to determine the relationship between nanoscale structural changes and the microscale modification observed in PNNs post-sensory deprivation and pathology, in particular. The X-ray diffraction method is also quicker than using IHC, reducing time sectioning, staining, and visualizing tissue. This could be extremely valuable when processing large sample quantities.

## Acknowledgements

This work was supported by NIGMS 122657 to JCB.

## Disclosures/Conflicts

JCB, EW are employees of the City University of New York (CUNY)

EW receives stipend support from the National Institutes of Health

S. M. is a paid intern at Matur UK, Ltd.

A.L. and P.L. are employed by Arion Diagnostics, Inc.

P.L. is employed by Matur UK, Ltd.

P.L is a shareholder of Matur UK, Ltd.

P.L. is a shareholder of Arion Diagnostics, Inc.

The authors declare no additional conflicts of interest.

## Notes

### Competing Interest Statement

JCB and EW are employees of the City University of New York (CUNY)
EW receives stipend support from the National Institutes of Health
S. M. is a paid intern at Matur UK, Ltd.
A.L. and P.L. are employed by Arion Diagnostics, Inc.
P.L. is employed by Matur UK, Ltd.
P.L is a shareholder of Matur UK, Ltd.
P.L. is a shareholder of Arion Diagnostics, Inc.
The authors declare no additional conflicts of interest.

### Summary of Updates

Provided additional clarifications as well as some additional analyses.

